# In *Staphylococcus aureus*, MbcS is a refunctionalized acyl-CoA synthetase that confers a fitness advantage during intra-species competition

**DOI:** 10.64898/2026.03.28.714991

**Authors:** Marcelle C. dos Santos Ferreira, Timothy G. Stephens, Shaun R. Brinsmade

**Affiliations:** Department of Biology, Georgetown University, Washington, DC, USA; Department of Biochemistry and Microbiology, School of Environmental and Biological Sciences, Rutgers University, New Brunswick, NJ, USA

**Keywords:** *Staphylococcus aureus*, acyl-CoA synthetase, branched-chain fatty acids, BKD complex, membrane biogenesis, competition

## Abstract

*Staphylococcus aureus* is one of the most frequently co-isolated pathogens in polymicrobial infections, where interspecies interactions contribute to enhanced virulence, persistence, and antimicrobial tolerance. Nutrient availability plays a central role in these interactions as microorganisms compete for resources required to sustain essential cellular processes. For instance, branched-chain amino acids (BCAAs) are critical for protein synthesis, and valine synthesis pathway precursors are essential for energy production. In *S. aureus*, BCAAs are also the precursors for branched-chain fatty acids (BCFAs), the dominant fatty acids in the *S. aureus* membrane. We previously identified a second pathway that uses branched-chain carboxylic acids (BCCAs) and the high-affinity acyl-CoA synthetase MbcS to catalyze the synthesis of BCFA precursors. However, the physiological role of this pathway and the conditions triggering its activation remain unclear. Here, we show that *mbcS* is restricted to *S. aureus* and closely related human-associated staphylococci. Phylogenetic analyses suggest that MbcS arose from a refunctionalization event and represents a non-orthologous replacement for the phosphotransbutyrylase (Ptb) and butyrate kinase (Buk) enzymes. Consistent with this model, Ptb and Buk from *Staphylococcus pseudintermedius* catalyze the formation of branched-chain acyl-CoAs from BCCAs, but only at high substrate concentrations. We further show that *mbcS* expression is upregulated in a *codY* mutant, implicating this pathway in BCAA-limited conditions. In support, we show that *mbcS* is required for optimal fitness during intra-species competition. Together, our findings support a model in which the MbcS-dependent pathway enables *S. aureus* to scavenge BCFA precursors under nutrient-limited conditions, providing a competitive advantage in polymicrobial environments.

**Importance:** *Staphylococcus aureus* is a major contributor to polymicrobial infections, where competition for nutrients can influence bacterial physiology and survival. A deeper understanding of how *S. aureus* adapts to nutrient limitation is therefore essential to explain its success as a human pathogen. In *S. aureus*, the acyl-CoA synthetase MbcS supports BCFA synthesis from BCAA-derived carboxylic acids and aldehydes, which are released into the environment as by-products of bacterial metabolism. Herein, we provide evidence that *S. aureus* acquired the acyl-CoA synthetase MbcS as an adaptive trait. This metabolic innovation allows this bacterium to maintain membrane homeostasis under nutrient limitation and compete against neighboring bacteria. Our findings highlight an adaptive strategy that may contribute to the persistence of *S. aureus* in polymicrobial infections.

## Introduction

*Staphylococcus aureus* is a commensal bacterium and an important opportunistic pathogen. The organism is the leading cause of skin and soft tissue infections, cutaneous abscesses, and a leading cause of life-threatening infections such as endocarditis, osteomyelitis, toxic shock syndrome, and bacteremia (1–4). *S. aureus* is also a major contributor to polymicrobial infections, such as those in chronic wounds and cystic fibrosis (5–8). In the polymicrobial environment synergistic interactions between microbial species enhance virulence, biofilm formation, persistence, and antibiotic tolerance compared with their mono-infection counterparts (5, 9, 10). Furthermore, organisms in a polymicrobial community face nutrient starvation, as they compete for the available nutrients to maintain vital metabolic processes (11, 12). Within the polymicrobial environment, *S. aureus* can establish either cooperative or competitive relationships with other microbial species (5). However, knowledge of the mechanisms that underly these synergistic interactions is still incomplete and further understanding of these mechanisms is critical for developing new therapies.

Amino acids, among other nutrients, are essential for growth and persistence of *S. aureus*, as they are involved in protein synthesis, membrane biogenesis, and regulation of virulence factors (13–15). Thus, amino acid starvation can impair vital cellular functions and prevent growth and survival. The global transcriptional regulator CodY monitors branched-chain amino acid (BCAA; i.e., isoleucine, leucine, valine) sufficiency as well as intracellular GTP levels and adjusts the expression of dozens of genes in a number of Gram-positive genera in response (16–20). In *S. aureus,* CodY controls both metabolism genes (i.e., genes for alternative nutrient uptake and processing, etc.) and virulence genes (14, 21–25). Notably, BCAAs also serve as precursors for the synthesis of BCFAs, the predominant fatty acids in the *S. aureus* membrane (26, 27).

BCFAs are essential for membrane fluidity, and to prevent membrane phase separation, protein segregation, and cell death. They are also essential for full virulence (25, 28–30). Their synthesis starts with the transamination of BCAAs by the aminotransferase IlvE. The α-keto acid products then undergo oxidative decarboxylation and subsequent activation catalyzed by the branched-chain α-keto acid dehydrogenase complex (BKDH) to generate acyl-CoA primers derived from isoleucine, leucine, and valine (2-methylbutyryl-CoA, 3-methylbutyryl-CoA, and isobutyryl-CoA, respectively). These primers are then elongated iteratively by the fatty acid synthase complex (i.e., FASII) to assemble long, branched-chain fatty acids (31, 32). In *S. aureus* the BKDH complex is composed of four enzymes: a dehydrogenase (E1α), a decarboxylase (E1β), a dihydrolipoamide acyltransferase (E2), and a dihydrolipoamide dehydrogenase (E3), encoded by *bkdA1*, *bkdA2*, *bkdB*, and *lpdA*, respectively (28). In other Gram-positive bacteria, such as *Bacillus subtilis* and *Enterococcus faecalis*, the *bkd* locus contains two additional genes: *ptb* and *buk*, encoding phosphotransbutyrylase and butyrate kinase, respectively (33, 34). During the catabolism of BCAAs in these organisms, Ptb transfers inorganic phosphate to the acyl-CoA substrate forming acyl-phosphate. Subsequently, the butyrate kinase Buk phosphorylates ADP, yielding ATP via substrate level phosphorylation and BCCAs. At least in *E. faecalis,* BCCA byproducts of catabolism are expelled into the environment (34, 35). Interestingly, biochemical characterization of Ptb and Buk from *Listeria monocytogenes* revealed that these enzymes are active in the reversible direction and can form BC-acyl-CoA precursors from BCCAs (36, 37). Whether this reversibility also occurs in vivo is yet to be studied.

Acetate is an abundant short-chain fatty acid and crucial for many metabolic processes in prokaryotes and eukaryotes, notably as source of acetyl groups for the tricarboxylic acid cycle and for straight-chain fatty acid synthesis. In bacteria, depending on carbon source availability, acetate can be assimilated or dissimilated. This change in metabolism is called the acetate switch (38). This has been studied extensively in the model organisms *Escherichia coli* and *Salmonella enterica.* In these bacteria during rapid exponential growth in the presence of the preferred carbon source glucose, the hexose is catabolized via the glycolytic pathway, generating pyruvate. Pyruvate is then oxidized to acetyl-CoA by the pyruvate dehydrogenase complex, analogous to the BKDH complex described above. When cellular levels of acetyl-CoA exceed demand, the acetyl-CoA is converted to acetate and excreted extracellularly by the coordinated action of the low-affinity enzymes phosphotransacetylase (Pta) and acetate kinase (Ack) (38–41). This allows the cells to make ATP, and regenerates coenzyme A for other biological processes. During post-exponential phase growth when glucose is exhausted, acetate is consumed and enters the tricarboxylic acid cycle via the high-affinity acetyl-CoA synthetase Acs (39–42). *S. aureus* carries out a similar “overflow” metabolism using Pta/AckA and AcsA (43–45). Notably, our group and others discovered *S. aureus* methylbutyryl-CoA synthetase (*Sa*MbcS) that is 56% similar and 37% identical end-to end to *S. aureus* acetyl-CoA synthetase AcsA, and 43%-47% similar to other bona fide acyl-CoA synthetases from *S. enterica* and *Rhodopseudomonas palustris*. Despite its structural and functional similarities to AcsA and other acyl-CoA synthetases, MbcS shows very low enzymatic activity with acetate. Instead, we found that its preferred substrates are the BCCAs isobutyric acid (*i*C_4_) and 2-metylbutyric acid (*a*C_5_) derived from isoleucine and valine, respectively (46). Isovaleric acid (*i*C_5_), derived from leucine, also serves as substrate, albeit poorly. MbcS can activate endogenous and exogenous BCCAs to generate acyl-CoA primers and promotes BCFAs synthesis in the absence of the canonical BKDH-dependent pathway (46, 47).

Herein, we extend our previous studies on MbcS. We report bioinformatics analysis revealing *mbcS* is restricted to *S. aureus* and related human-associated staphylococcal genomes. Our data suggest that MbcS originated from a refunctionalization event and is a non-orthologous replacement for Ptb and Buk. In support, we found that Ptb and Buk from the canine commensal and opportunistic pathogen *S. pseudintermedius* can catalyze the formation of branched-chain acyl-CoA primers from short BCCAs. The high BCCAs levels required for *Sp*Buk activity suggest that Ptb and Buk comprise a low-affinity biosynthetic pathway. We demonstrate that *mbcS* is upregulated in a *codY* null mutant, which led us to hypothesize that the MbcS-dependent pathway is important in a BCAA-limited environment. Notably, although *mbcS* is dispensable for *S. aureus* growth in monoculture, MbcS-deficient cells exhibit a fitness disadvantage relative to MbcS-proficient cells in direct competition assays. Our studies bring a greater understanding of the role of MbcS in *S. aureus* physiology. In particular, we posit that MbcS plays an important role in the polymicrobial environment.

## Results

### The acyl-CoA synthetase MbcS is only found in well-established human-associated staphylococci

We and others previously demonstrated that the *S. aureus* gene with locus tag SAUSA300_2542 encodes the acyl-CoA synthetase MbcS (46, 47). *mbcS* is found in the genome of USA300 as shown in **Fig. 1A**. To determine the extent that *mbcS* is conserved in bacteria, we performed a gene neighborhood analysis using the web server AnnoView (48) and found that *mbcS* is only present in *S. aureus* and other human-associated staphylococci such as *S. epidermidis*, *S. haemolyticus*, *S. hominis*, *S. capitis*, and *S. warneri*. Notably, these species are members of the human microbiota and opportunistic pathogens (49–53). Interestingly, our neighborhood analysis revealed *mbcS* is absent in other Gram-positive species including *Bacillus subtilis, Enterococcus faecalis, Listeria monocytogenes,* and the canine skin commensal and pathogen *S. pseudintermedius* (54, 55). Instead, these species harbor genes encoding putative phosphotransbutyrylase (*ptb*) and butyrate kinase (*buk*) in their *bkd* operons (**Fig. 1B**). In principle, these enzymes reversibly catalyze acyl-CoA formation from carboxylic acids. MbcS, Ptb, and Buk are reminiscent of high- and low-affinity Acs and Ack-Pta pathways for acetate consumption (41). In support, Sirobhushanam *et al.* provide biochemical evidence that *Listeria monocytogenes* Ptb and Buk catalyze acyl-CoA formation. Therefore, we hypothesized that *mbcS* is a non-orthologous replacement for *ptb* and *buk* in human-associated staphylococci. To start to test this hypothesis, we performed a phylogenetic analysis of all proteins annotated in NCBI as “Acyl-CoA synthetase”, “AMP-binding enzyme”, “MbcS”, or “Acs”. We identified at least three families of proteins which were all shared in the common ancestor of *Staphylococcus* (see blue, green, and yellow outer rings in **Fig. S1**). Only one of these protein families (Family 2 [green outer line] in **Fig. S1**) contained all putative MbcS sequences identified using their proximity in the genome to *betA* and *betB* (our high-confidence syntenicly-identified MbcS proteins; see *Material and Methods*). That is, while there are multiple families identified, only one appears to exclusively contain our high-confidence MbcS proteins. While only some of the proteins in the putative MbcS family were identified by our synteny analysis, we propose that all proteins in this clade (Family 2 in **Fig. S1**) are MbcS, with many escaping our initial search strategy due to variation in their functional descriptions. This analysis also suggests that homologs of the MbcS proteins are present in the outgroup species. While we do not suggest that these homologous outgroup proteins function as MbcS, we do propose that MbcS evolved from a refunctionalization of an existing protein family, rather than through gene duplication. Additionally, whereas many of the proteins in our putative *mbcS* gene clade were not identified by syntenic analysis, phylogenetic analysis of each genome used in our study demonstrated clear taxonomic restriction of the MbcS family proteins to human-associated *Staphylococcus* species (**Fig. 2**). This further strengthens the assertion that MbcS evolved as an adaptation to a human-associated lifestyle. Additionally, the phylogenetic analysis strengthened our previous neighborhood analysis as it shows that *ptb* and *buk* are found in the outgroup species and mostly in the non-human associated staphylococci, but not in *S. aureus* and other well-established human-associated species (**Fig. 2**).

**Figure 1.**
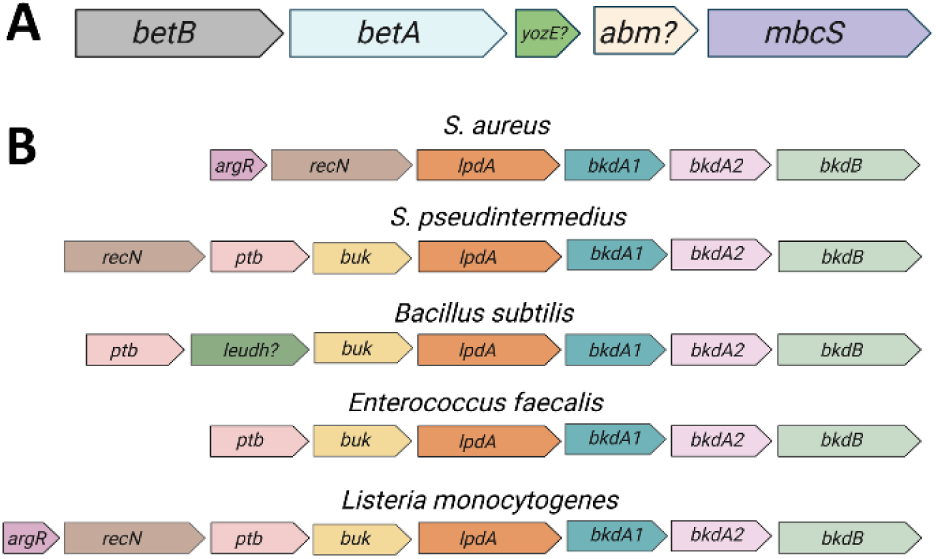
*mbcS* is not conserved in Gram-positive bacteria. **A.** Neighborhood analysis shows the genetic context of *mbcS.* The gene is found only in *S. aureus, S. epidermidis, S. haemolyticus, S. lugdunensis, S. hominis, S. capitis,* and *S. warneri*. *mbcS* is flanked by *betB* (glycine betaine aldehyde dehydrogenase), *betA* (choline dehydrogenase), and two genes coding for hypothetical proteins *yozE* (YozE family protein) and *abm* (antibiotic biosynthesis monooxygenase family protein). **B.** The *bkd* loci in selected Gram-positive bacteria. These bacteria harbor *ptb* and *buk,* but not *mbcS*. *argR* (arginine repressor), *recN* (DNA repair protein RecN), *ptb* (phosphotransbutyrylase), *buk* (butyrate kinase), *lpdA* (dihydrolipoamide dehydrogenase), *bkdA1* (2-oxoisovalerate dehydrogenase, α-subunit), *bkdA2* (2-oxoisovalerate dehydrogenase, β-subunit), *bkdB* (dihydrolipoamide acetyltransferase), *leudh* (leucine dehydrogenase).

**Figure 2.**
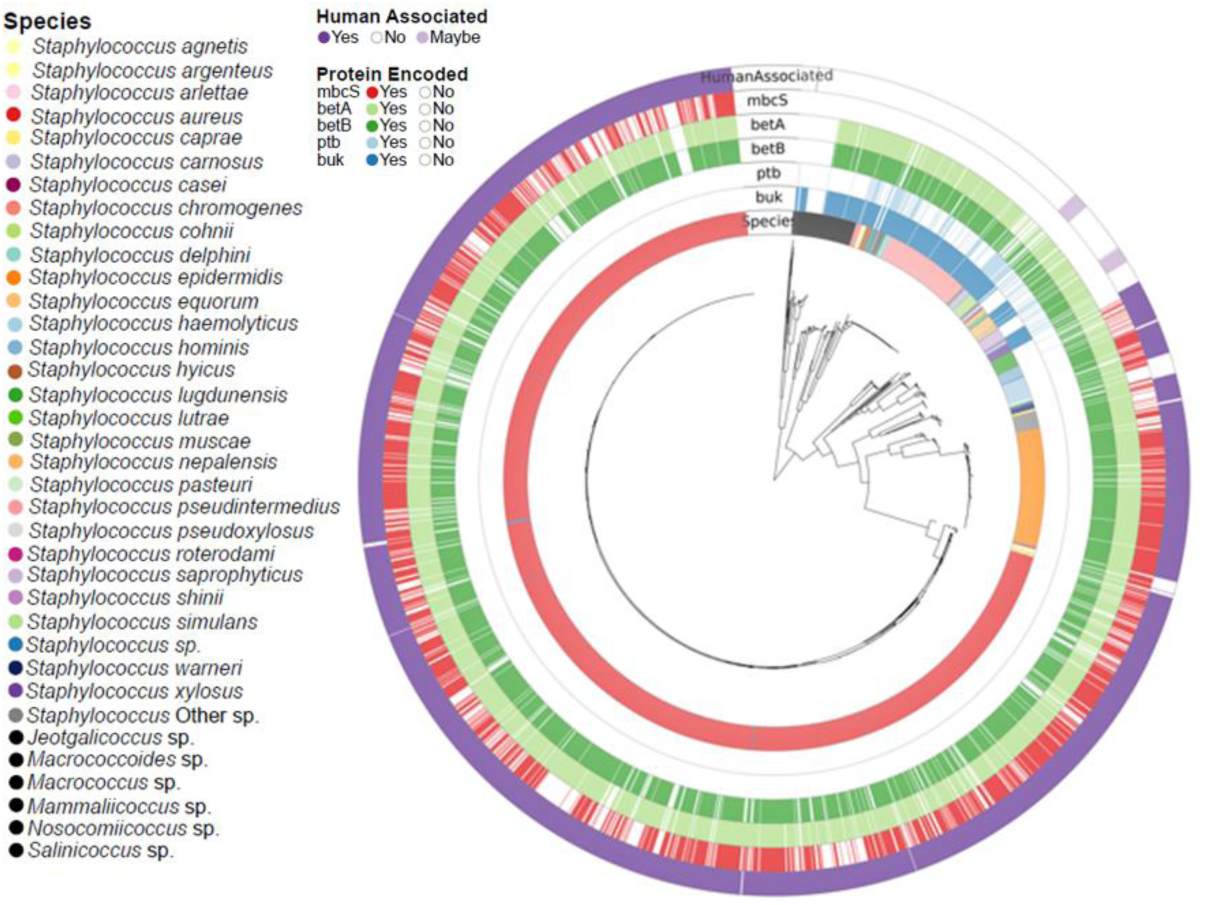
Occurrence of *mbcS* coincides with loss of *ptb* and *buk* in *S. aureus* and other human-associated staphylococci. Phylogenetic analysis was performed taking into account the genetic context of *mbcS* and presence and absence of *ptb* and *buk* in *Staphylococcus species*. Assembled Staphylococcaceae genomes with predicted proteins were downloaded from the NCBI database.

### A putative acyl-CoA synthetase from *S. simulans* can promote *S. aureus* growth from exogenous short BCCAs

Curiously, while proteins that fall into the Family 2 clade are phylogenetically MbcS, we found some MbcS-coding genes lack genetic synteny. For instance, the *S. simulans* strain IVB6244 genome (GenBank accession number CP094697) contains the gene locus MUA81_01165 annotated to encode the AMP-binding protein UXR33001.1, which is 81% similar and 69% identical (end-to-end) to *Sa*MbcS. Furthermore, this strain has *ptb* and *buk* as part of the *bkd* operon. Interestingly, besides their high sequence similarity, *Sa*MbcS and UXR33001.1 also share the C-terminal catalytic lysine 510 residue (K510), conserved among this family of acyl-CoA synthetases (**Fig. 3A**) (39). This suggests that UXR33001.1 has acyl-CoA synthetase activity and may catalyze the conversion of exogenous BCCAs to branched-chain (BC) acyl-CoAs. To test this genetically, we cloned the MUA81_01165 gene from *S. simulans* into the integrative, single-copy plasmid pCT3 (46), which places the MUA81_01165 gene under the control of the inducible P*_tet_* promoter. We then introduced this plasmid into the *S. aureus lpdA::kan^+^ mbcS::erm^+^* double mutant (a BCFA auxotroph) and assessed the ability of the resulting strain to grow when fed BCCAs exogenously in chemically defined medium (CDM). As expected, the double mutant exhibited BCFA auxotrophy and failed to grow beyond an optical density at 600 nm (OD_600_) of ∼0.1 when BCCAs were not provided, and auxotrophy was suppressed when *SambcS^+^* and a mixture of all three BCCAs were provided (**Fig 3B-C**, compare green vs. gray bars). Importantly, ectopic expression of MUA81_01165 also suppressed auxotrophy, albeit to a lesser degree than *SambcS^+^*(**Fig. 3B-C**). We note that inducing expression with 10 ng ml^-1^ anhydrotetracycline (aTc) and increasing the BCCA concentration to 1 mM did not enhance growth under the same growth conditions. We cannot exclude the possibility that protein levels are insufficient to support wild-type culture yields as our MbcS antibodies do not cross-react with UXR33001.1 (**Figs 3C, S2, and data not shown**). Nevertheless, our data suggest that UXR33001.1 can catalyze BC-acyl-CoA synthesis.

**Figure 3.**
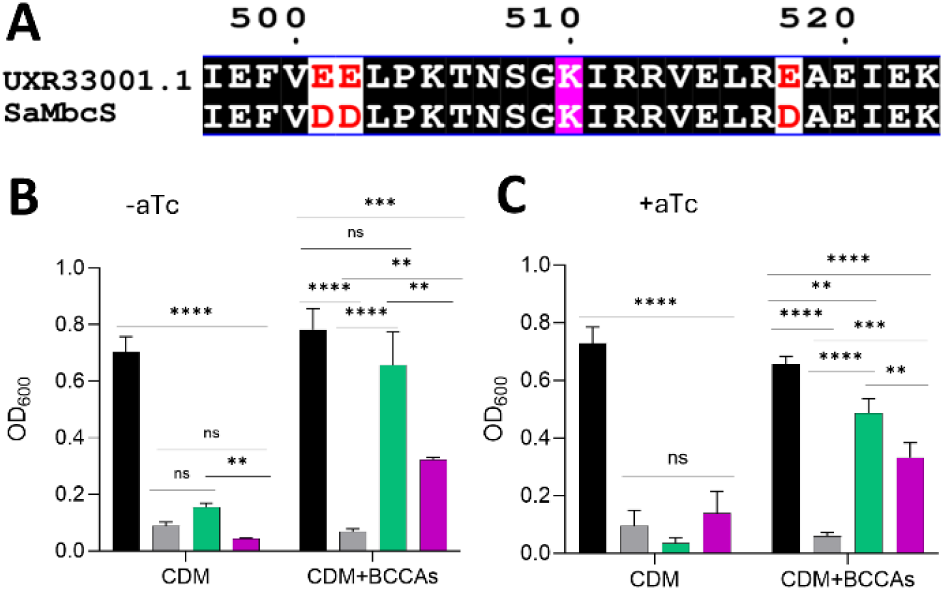
UXR33001.1 is a putative acyl-CoA synthetase from *S. simulans*. (**A**). ClustalW (98) was used to align MbcS from *S. aureus* and the putative acyl-CoA synthetase UXR33001.1 from *S. simulans*. Sequences were formatted using ESPript (99). A C-terminal portion of the sequence alignment is shown with amino acids strictly conserved between the proteins shaded; the conserved lysine residue required for the catalytic activity of acyl-CoA synthetases is highlighted in magenta. (**B,C**). WT harboring pWY53 (pCT3 empty vector; black bar) and the *lpdA::kan^+^ mbcS::erm^+^* double mutant harboring either pWY53 (grey bar), pWY54 (pCT3 with *SambcS*^+^; green bar), or pMF022 (pCT3 with MUA81-01165^+^; purple bar) were grown in CDM or in CDM supplemented **(B)** with 500 µM BCCAs with or **(C)** without anhydrotetracycline (aTc). Optical density at 600 nm (OD_600_) was measured after 20 h. Data are plotted as mean ± SD from three biological replicates. *\*\*\*\**p<0.0001, ***p<0.001, **p<0.01, one-way ANOVA with Tukey’s multiple comparison test; ns, not significant.

### The Ptb-Buk pathway from *S. pseudintermedius* shows low activity toward BCCAs in laboratory culture

Biochemical characterization of Pta-AckA and Buk-Ptb suggest these enzymes comprise low-affinity pathways, as they demonstrate high *K_m_*values (i.e. in the millimolar range) for acetate and BCCAs, respectively (36, 37, 39, 41). Conversely, we previously demonstrated that, like Acs in *Salmonella enterica*, MbcS is a high-affinity acyl-CoA synthetase with *K_m_* values ranging from 5 to 10 µM for the BCCAs derived from isoleucine (*a*C_5_) and valine (*i*C_4_), respectively. It can also utilize the leucine derivative (*i*C_5_) when its concentration is above 500 µM (46). Based on our phylogenetic findings that presence of *ptb* and *buk* is anticorrelated with *mbcS* in the staphylococci genome, we hypothesize that Ptb and Buk comprise a low-affinity pathway (**Fig 4A**), which was replaced by MbcS in *S. aureus*. If so, MbcS would allow human-associated staphylococci to utilize even very low quantities of BCCAs available in the infection niche to generate BC-acyl-CoAs.

**Figure 4.**
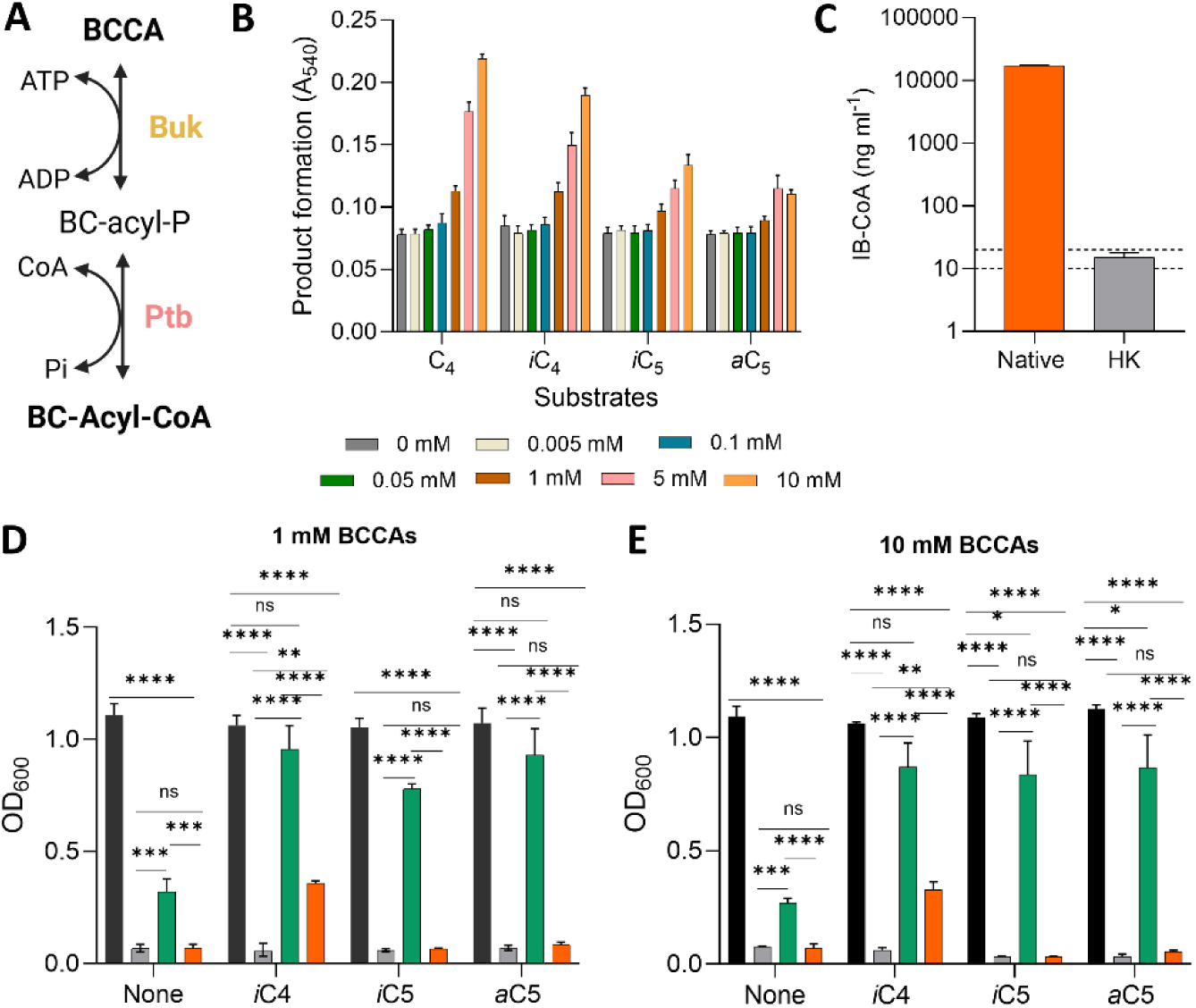
*ptb^+^*and *buk^+^* from *S. pseudintermedius* partially restore growth of the *S. aureus lpdA::kan^+^ mbcS::erm^+^* double mutant in CDM supplemented with *i*C_4_. (**A**) Proposed pathway for the conversion of BCCAs to acyl-CoA derivatives via Buk and Ptb. (**B**) The activity of *Sp*Buk was tested in vitro using the hydroxamate assay as described in the *Materials and Methods* with various concentrations of short, straight and branched carboxylic acids. Acids are indicated as C*_X_*, where *X* denotes the carbon length. Data are plotted as the mean absorbance at 540 nm (A_540_) for acyl-hydroxamate formation ± SD after 1 h of three independent trials. (**C**) Native or heat killed (HK) *Sp*Ptb and *Sp*Buk were incubated with MgCl_2_, ATP, TCEP, CoA, and 10 mM (2 µmol) of isobutyric acid (IB) to produce IB-CoA as described in *Materials and Methods*. Detection and quantification of IB-CoA was measured via LC–MS. Reactions with HK enzymes yielded very small amounts of IB-CoA that were above the limit of detection (10 ng ml^-1^) but below the lower limit of quantification (20 ng ml^-1^) (gray bar and dashed lines). Data are representative of three independent trials. (**D-E**) WT cells harboring pWY53 (pCT3 empty vector; black bar) and *lpdA::kan^+^ mbcS::erm^+^* double mutant cells harboring either pWY53 (grey bar), pWY54 (pCT3 with *SambcS*^+^; green bar), or pMF020 (pCT3 with S*pptb*^+^*buk*^+^; orange bar) were grown in CDM or CDM supplemented with either (**D**) 1 mM or (**E**) 10 mM of the respective BCCAs, without anhydrotetracycline as inducer. Optical density at 600 nm (OD_600_) was measured after 20 h. Data are plotted as mean ± SD from three biological replicates. *\*\*\*\**p<0.0001, ***p<0.001, **p<0.01, *p<0.05, one-way ANOVA with Tukey’s multiple comparison test within each condition; ns, not significant.

To characterize Buk and Ptb directly, we overproduced Buk and Ptb from *S. pseudintermedius (Sp*Buk and *Sp*Ptb) in *E. coli* BL21(DE3). The enzymes were purified to near apparent homogeneity using an N-terminal hexahistidine tag and Ni^2+^-NTA chromatography. Purified *Sp*Buk was tested for branched-chain carboxylic acid kinase activity by incubating the protein with Mg*ATP and various fatty acid substrates and monitoring the formation of the acyl-phosphate product using the hydroxamate assay (56, 57). Our data demonstrate that *Sp*Buk exhibits the highest activity toward butyric acid (C_4_). The enzyme was also capable of catalyzing the formation of acyl-P from *i*C_4_ as well as with *a*C_5_ and *i*C_5_, though we observed a clear preference for C_4_ substrates (**Fig. 4B**). No product was formed when *Sp*Buk protein was heat-inactivated before incubation with the substrates (**data not shown)**. Importantly, *Sp*Buk only shows substantial activity with high substrate concentrations (i.e. 5 mM and 10 mm) (**Fig. 4B**). Next, to test if the Buk-Ptb pathway is active in the acyl-CoA forming direction, we incubated recombinant *Sp*Buk and *Sp*Ptb with 10 mM *i*C_4_, Mg*ATP, and coenzyme A. Isobutyryl-CoA (IB-CoA) formation was quantified by mass spectrometry (LC-MS) analysis. As demonstrated in **Fig. 4C**, IB-CoA was detected in reactions containing active enzymes, but virtually no IB-CoA was detected when enzymes were heat-inactivated prior to incubation with substrates.

To lend support for the hypothesis that Buk-Ptb comprise a low-affinity pathway to generate BC acyl-CoA primers for BCFAs synthesis, we assessed the ability of *Spptb^+^* and *Spbuk^+^* to complement an *S. aureus* strain defective in BC acyl-CoA formation. We used the *S. aureus lpdA::kan^+^ mbcS::erm^+^* double mutant to test *Spbuk^+^* and *Spptb^+^* function. Plasmid pMF020 (*Spbuk^+^*, *Spptb^+^*) was introduced into this strain, along with pCT3 as the negative control and pWY54 (*SambcS^+^*) as the positive control. Growth behavior was assessed in either chemically defined medium (CDM) or CDM supplemented with 1 mM or 10 mM of the individual BCCAs (**Fig. 4D-E**). In the presence of 1 mM of the individual BCCAs, ectopic expression of *SambcS^+^* in the *lpdA::kan^+^ mbcS::erm^+^* double mutant suppressed branched-chain fatty acid auxotrophy essentially to WT levels with all three BCCAs (**Fig. 4D**). Consistent with our in vitro data showing relatively efficient conversion of *i*C_4_ to acyl-P and IB-CoA, complementation with *Spbuk^+^-ptb^+^* promotes partial growth of the *lpdA::kan^+^ mbcS::erm^+^* strain in the presence of *i*C_4_; we detected essentially no growth when *a*C_5_ *and i*C_5_ were provided (**Fig. 4D**). Interestingly, no further growth enhancement was observed when we increased the concentration of each BCCA to 10 mM or induced expression with aTc (**Figs. 4E, S3**), suggesting that activity of the Ptb-Buk pathway in the acyl-CoA formation direction is limited in vivo. However, we cannot exclude the possibility that the poor growth observed is due to suboptimal protein production. Our efforts to complement the *lpdA::kan^+^ mbcS::erm^+^* double mutant with epitope-tagged proteins failed and antibodies against *Sp*Ptb and *Sp*Buk are not available. Interestingly, we observed mild growth of the *lpdA::kan^+^ mbcS::erm^+^*double mutant with *SambcS*^+^ in CDM without exogenous BCCAs (**Fig. 4D-E**). This reinforces our previous observation that MbcS can also generate BC acyl-CoA primers from internal carboxylic acids pools produced by yet to-be-identified enzymes (46).

### *mbcS* is upregulated when CodY activity is reduced

In our previous work we identified the acyl-CoA synthetase MbcS in a *lpdA::erm^+^* mutant background, where the BKDH-dependent pathway is inactive. In that study, we found promoter-up mutations at the *mbcS* locus that restored BCFA synthesis (46). Why does *S. aureus* have two pathways for BCFA synthesis? Is *mbcS* ever upregulated? Which physiological conditions might upregulate *mbcS* transcription and trigger the MbcS-dependent pathway? To begin to answer these questions, we screened a subsection of the Nebraska Transposon Mutant Library (NTML) using a P*_mbcS_-gfp* transcriptional reporter plasmid (58, 59). Each mutant contains a disruption in one of the 111 known and putative transcription factors in *S. aureus.* Because we do not yet know the conditions (if any) that induce expression of *mbcS^+^*, our approach is designed to find transcriptional repressors. pAP4 (P*_mbcS_-gfp*) was transduced into the library creating 111 strains simultaneously. The strains were then screened in chemically defined medium supplemented with 10 g L^-1^ tryptone, allowing us to better discern any increase in *mbcS^+^* promoter activity. Three of the 111 transposon mutants - *codY::erm^+^, srrA:erm^+^,* and an unknown putative transcriptional regulator (SAUSA300_0577) - showed increased promoter activity compared with the wild-type strain (**Fig. S4**). The SAUSA300_0577 gene product likely controls the expression of the downstream gene, the putative pyridine nucleotide-disulfide oxidoreductase (SAUSA300_0576). As such, this presumptive “local” regulator was not pursued further. We then focused on the two known global transcriptional regulators SrrA and CodY. After reconstructing strains and remeasuring promoter activity, the *codY::tet^+^* mutant exhibited a reproducible and significant two-fold increase in P*_mbcS_-gfp* promoter activity compared with the WT parent strain (**Fig. 5A-B**). Importantly, the increased promoter activity phenotype in the *codY::tet*^+^ mutant was suppressed when we introduced a wild-type copy of *SacodY*^+^ (**Fig. 5B**).

**Figure 5.**
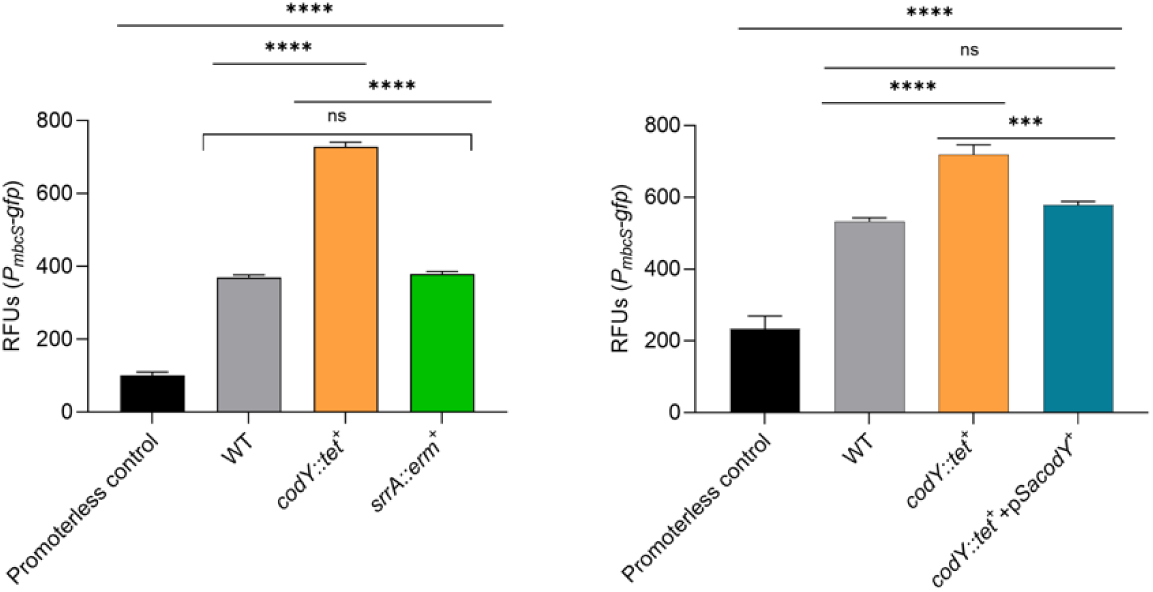
*mbcS* is overexpressed in a *codY* mutant background. (**A**) *S. aureus* WT, *codY::tet^+^*, and *srrA::erm^+^* mutant cells with plasmid pAP4 containing the *mbcS^+^* promoter region fused translationally to *gfp* (P*_mbcS_-gfp*) were grown to stationary phase in CDM supplemented with 10 g L^-1^ tryptone. Cells were pelleted and washed with PBS. Promoter activity was then measured (relative fluorescence units [RFUs]; GFP/OD_600_). (**B**) WT, *codY::tet^+^*, or *codY::tet^+^* cells complemented with p*SacodY*^+^ harboring P*_mbcS_-gfp* were grown to stationary phase in TSB, at which time cells were pelleted and washed with PBS. Promoter activity was then measured (RFUs; GFP/OD_600_). In both cases the WT strain harboring promoterless plasmid was used as a negative control. *\*\*\*\**p<0.0001 one-way ANOVA with Tukey’s multiple comparison test. ns, not significant. n=3.

### MbcS gives *S. aureus* a competitive advantage in a nutrient-limited environment

In line with the role of CodY in prioritizing gene expression as a function of nutrient sufficiency (22, 60), we wondered if, under low BCAA availability, *S. aureus* prefers to utilize exogenous branched-chain carboxylic acids for membrane biogenesis via MbcS instead of using BCAAs for BCFAs synthesis through the canonical BKDH-dependent pathway. During infection, intra- and interspecies competition for BCAAs would likely be severe. We hypothesized that cells expressing *mbcs^+^* would give *S. aureus* a competitive advantage under these conditions. To test this hypothesis, we performed a competition assay to assess fitness conferred by *mbcS^+^* during growth in rich, complex medium (TSB). *S. aureus* WT cells marked with trimethoprim or tetracycline resistance (SRB1710 and SRB3143, respectively), *mbcS::erm^+^*mutant cells carrying pCT3 (marked with erythromycin and tetracycline resistance; SRB3107) and *mbcS::erm^+^*mutant cells complemented with p*SambcS*^+^ (marked with erythromycin resistance and tetracycline resistance; SRB3108) were grown either as monocultures or as co-cultures as described in *Material and Methods*. Drug resistance markers allowed us to distinguish strains by counting colony forming units (CFUs) on selective media. For cells grown as monocultures, there were no differences between strains either in initial (input) or in final (output) CFU counts (**Fig. S5A-B**). In co-cultures, initial CFU counts were roughly the same and no significant differences were observed between strains (**Fig. S5C**). However, when WT cells were co-cultured with *mbcS::erm^+^* cells, we observed a significant decrease in the final CFU counts in the latter strain (**Fig. S5D**). We measured no difference in CFU counts when wild-type cells marked with different antibiotic resistance genes were competed 1:1 or when WT cells were competed with *mbcS::erm^+^* cells complemented with p*SambcS*^+^. We then calculated the competitive index as previously described (61, 62). As expected, there was no difference in fitness in WT cells carrying different drug markers (**Fig. 6**, grey symbols). However, we calculated a 10-fold decrease in the competition index when *mbcS::erm^+^*cells were co-cultured with wild-type cells, indicating that MbcS provides a fitness advantage. This fitness cost was abrogated when we complemented the *mbcS::erm*^+^ mutant with p*SambcS^+^* (**Fig. 6**, compare blue and pink symbols). These data suggest that, as nutrients become limited due to competition, the ability to use exogenous BCCAs for BCFAs synthesis via the MbcS-dependent pathway provides a fitness advantage.

**Figure 6.**
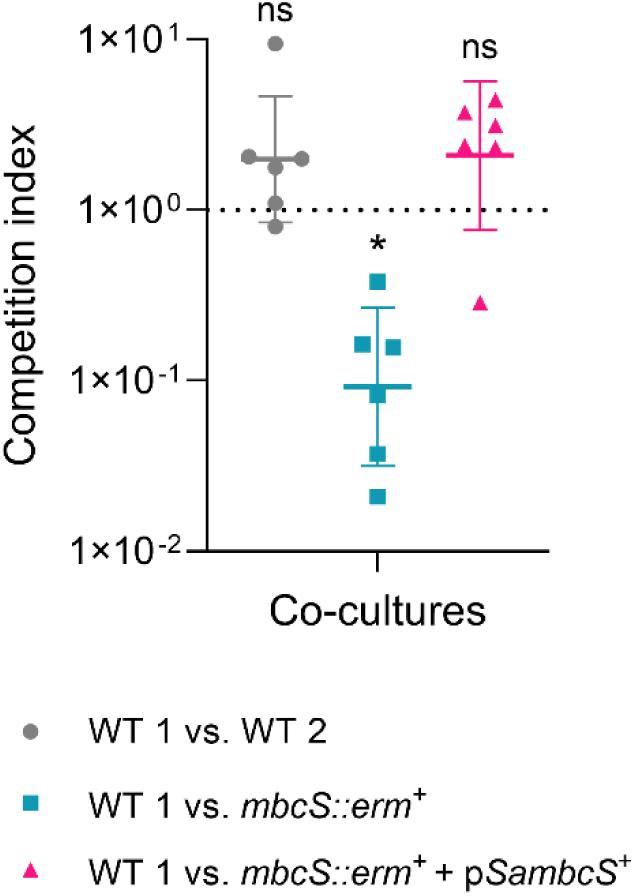
*mbcS^+^*confers a fitness advantage during intra-species competition. *S. aureus* CFU counts from the three independent co-cultures (see **Fig S5**) were used to calculate the respective competition indices (CIs). Mutant/WT output ratios were divided by Mutant/WT input ratios to determine CIs (61, 62). Dashed line at 1X10^0^ denotes CI=1 and no fitness phenotype. Data are plotted as mean ± SD from six biological replicates within each co-culture. *p<0.05, one sample Wilcoxon test, where the null hypothesis is 1 (i.e. no competition); ns, not significant.

## Discussion

Branched-chain fatty acids (BCFAs) are the major fatty acids in the *S. aureus* membrane (26, 27). They are crucial for cell homeostasis, growth, stress resistance, virulence, and ultimately survival and persistence (25, 28, 29, 63). The critical role of BCFAs is highlighted by the presence of two pathways for their synthesis in *S. aureus*. We previously demonstrated that the MbcS-dependent pathway is upregulated and promotes BCFAs synthesis in vitro when the canonical BKDH-dependent pathway is disrupted (46). However, we wondered why *S. aureus* has acquired an additional pathway for BCFAs and what its physiological role would be. In the present study, we found that the *mbcS* gene is restricted to *S. aureus* and other well-established human associated staphylococci (**Figs. 1 and 2**). Our phylogenetic data provide strong evidence that *mbcS* is a non-orthologous replacement for the phosphotransbutyrylase *ptb* and the butyrate kinase *buk* (**Fig. 2**). We found that the putative acyl-CoA synthetase from *S. simulans* promotes bacterial growth in the presence of exogenous short branched-chain carboxylic acids (BCCAs) (**Fig. 3**). Additionally, the evidence that Ptb and Buk from *S. pseudintermedius* catalyzes BC acyl-CoA formation albeit from very high levels of short BCCAs unlikely to be present during infection suggests that *S. aureus* and other human-associated staphylococci evolved to use MbcS to scavenge BCFA precursors (**Fig. 4**). In support, we show that *mbcS* is dispensable for *S. aureus* growth in monocultures but is required for optimal fitness when in co-cultures. We have strong evidence that the MbcS-dependent pathway contributes to the synthesis of acyl-CoA primers from exogenous BCCAs when isoleucine, leucine, and valine are limited (**Figs. 5-6**).

In nature, duplicated genes can undergo subfunctionalization with specialization of the ancestral functions or they can acquire a new function, a phenomenon called neofunctionalization (64–66). We found that the *mbcS* gene is primarily organized in a specific genetic context, which is conserved only in a few human associated staphylococci (**Figs. 1 and 2**). *S. simulans* is a common zoonotic pathogen and is an emerging human opportunistic pathogen (67). Interestingly, our phylogenetic analysis discovered that the *S. simulans* gene locus MUA81_01165 is annotated to encode the putative acyl-CoA synthetase UXR33001.1, which is 81% similar and 69% identical end-to-end to *Sa*MbcS. Furthermore, this strain and other *S. simulans* strains have *ptb* and buk in their genomes (**Fig. 2**). Despite not being found in the same genetic context of *SambcS*, our data indicate that UXR33001.1 has acyl-CoA synthetase activity in vivo, as MUA81_01165 partially restores growth of the *lpdA::kan^+^ mbcS::erm^+^* double mutant in the presence of exogenous BCCAs (**Figs. 3 and S2**). Could MUA81_01165 be the ancestral gene that originated *mbcS* after a subfunctionalization event? Further investigation is needed to determine whether human-associated staphylococci refunctionalized MbcS as an adaptative trait. We know that the BCCAs *i*C_4_ and *a*C_5_ are the preferable substrates of *Sa*MbcS (46) (**Figs. 4 and S2**). However, our data suggest that UXR33001.1 does not have the same substrate specificity observed for *Sa*MbcS, as *lpdA::kan^+^ mbcS::erm^+^* with MUA81_01165^+^ grows to same extent in the presence of all three different BCCAs (**Fig. S2**). A biochemical characterization of UXR33001.1 would be required to support the observed growth phenotype.

Volatile compounds such as aldehydes and carboxylic acids can be synthesized and released for many bacterial species as byproducts of branched-chain amino acid catabolism (34, 68–71). These volatile compounds are important for flavor formation in the food industry and for targeted diagnosis of bacterial infections (71, 72). We previously described that the MbcS-dependent pathway forms BC-acyl-CoAs from exogenous BCCAs and from BCCAs generated from BC-aldehydes precursors. We proposed that α-keto acids are converted into their respective branched-chain aldehydes by an α-keto acid decarboxylase, followed by a reaction catalyzed by an aldehyde dehydrogenase to generate BCCAs (46). In support, we observed a slightly, yet significant, growth of *lpdA::kan^+^ mbcS::erm^+^* double mutant with *SambcS*^+^ in CDM without carboxylic acids (**Figs. 4 and S3**). The α-keto acid decarboxylase and the aldehyde dehydrogenase involved in the MbcS-dependent pathway are still to be found but this is currently an active focus of our laboratory.

The butyrate kinase Buk and the phosphotransbutyrylase Ptb are well known for mediating energy production in fermenting bacteria as they catalyze the conversion of an acyl-CoA to a carboxylic acid and concomitant formation of ATP (33–35, 73, 74). Ptb and Buk are analogous to Pta and Ack, which catalyze the reversible conversion of pyruvate to acetate under conditions of high intracellular acetate levels (i.e., a low-affinity pathway) (35, 39, 41). Here we show genetically and biochemically that Ptb and Buk from the canine-associated species *S. pseudintermedius* are active in the acyl-CoA forming direction (**Fig. 4A-C**). Buk activity increases with the increase in substrate concentration in the millimolar range, especially *i*C_4_ (**Fig. 4B**). These data indicate that these enzymes have a low affinity for their substrates. Our data are in strong agreement with the work of Sirobhushanam and colleagues, as they previously demonstrated that, in vitro, Ptb and Buk from *L. monocytogenes* can generate branched-chain acyl-CoA primers for BCFAs synthesis from high concentrations of exogenous BCCAs (36, 37). Is it possible that *S. aureus* and other human associated staphylococci gained MbcS functionality because running Ptb and Buk in the acyl-CoA forming direction is inefficient for the low quantities they would normally encounter? We observed activity of *Sp*Ptb*-Sp*Buk in vivo, as ectopic expression of *Spptb^+^-Spbuk^+^* partially restored growth of the *lpdA::kan^+^ mbcS::erm^+^* double mutant in CDM supplemented with *i*C_4_ (**Figures 4D-E, S3**). We posit that Ptb and Buk were subsequently lost to avoid futile cycling of key metabolites including ATP.

Amino acids are key precursors in many central metabolic pathways, including protein and nucleotide biosynthesis, which are vital for cellular processes across all kingdoms of life. The branched-chain amino acids (BCAAs) isoleucine, leucine, and valine serve as precursors for BCFAs synthesis in Gram-positive bacteria and the corepressors of the global transcriptional regulator CodY (14, 15, 20). Importantly, despite the presence of a genetic machinery for de novo synthesis of BCCAs, *S. aureus* typically relies on the uptake of exogenous BCAAs, which are transported to the cytoplasm via dedicated active transporters, named BrnQ1/2, and BcaP. This likely allows the bacterium to sense external ILV levels as a host cue to adjust metabolism and virulence (75–77). After screening a transcription factor mutant library for putative repressors of *mbcS* transcription (**Fig. S4**), we found that *mbcS* is significantly upregulated in the *codY* mutant background (**Fig. 5**). CodY’s activity is controlled by the levels ILV and GTP available. As these nutrients become limited, *S. aureus* CodY activity is reduced and its downstream targets derepressed (14, 22). It is worth noting that *mbcS* has not been identified previously as differentially expressed in *codY* mutant cells relative to wild-type cells, nor does the *mbcS* promoter appear to have an obvious CodY binding site (78). Nevertheless, *mbcS* promoter activity is increased when BCAA levels are perceived to be low. This mechanism would allow available BCAAs to be redirected to other vital cellular processes such as protein synthesis and pantothenate biosynthesis (79, 80). We suspect that CodY-dependent regulation of *mbcS* is indirect, and likely the result of subtle changes to metabolism. Does MbcS support *S. aureus* growth in a nutrient-limited environment, when the BKDH pathway is intact? Our data show that the *S. aureus mbcS::erm^+^* mutant struggles to grow in rich, complex medium (TSB) when in co-culture with wild-type parent strain, but not in monoculture (**Fig. S5B and D**). Additionally, having both the BKDH- and MbcS-dependent pathways active gives *S. aureus* a competitive advantage over the strain lacking a functional MbcS-dependent dependent pathway, also in TSB (**Fig. 6**). These findings are in line with the study of Qiao and colleagues, where they demonstrated that resource availability is a pivotal parameter for determining interactions within the community (12). Given the broad and vital roles isoleucine, leucine, and valine play in cellular functions (81), one might assume these would be the object of competition within a community, and would eventually become scarce. The high affinity of MbcS for short BCCAs substrates would compensate for the low BCAAs availability and preserve cell homeostasis.

We previously demonstrated that *mbcS* is heavily downregulated in our laboratory conditions. The spacing between the -35 and -10 regions in the *mbcS* promoter is not optimal and decreasing the spacing leads to upregulation of *mbcS* transcription (46). As such, strong transcriptional repression seems unnecessary. It is possible that a yet-to-be-identified positive regulator stimulates *mbcS* expression under certain conditions. Our laboratory is actively working to identify this regulator. In addition to transcriptional regulation, acyl-CoA synthetases are also subjected to post-translational modification as a way to regulate their enzymatic activity (82, 83). For instance, in *S. enterica*, Acs activity is regulated by reversible lysine acetylation mediated by the acetyltransferase Pat and the deacetylase CobB (84–86). Notably, acyl-CoA synthetases share a conserved C-terminus lysine residue required for activity, and that is part of the PX_4_GK acetylation motif (87, 88). Importantly, we demonstrated that the PX_4_GK acetylation motif is also conserved in both *Sa*MbcS and the putative acyl-CoA synthetase UXR33001.1 from *S. simulans* (46)(**Figure 3**). This opens up the possibility that *Sa*MbcS activity may be controlled by acylation. Experiments are ongoing in our laboratory to test this possibility.

As highlighted above, the catabolism of branched-chain amino acids can result in the synthesis of volatile compounds, including branched-chain carboxylic acids (BCCAs), that are subsequently released into the environment. Notably, *S. aureus* is commonly found in polymicrobial infections with some of the BCCAs-producers, such as *E. faecalis* and *Pseudomonas aeruginosa*. Specific niches where these organisms can be found together are chronic wounds and cystic fibrosis-related infections, respectively (7, 8). These observations along with our data presented herein informs our working model (**Fig 7**). During conditions where nutrients are freely available, *S. aureus* maintains membrane biogenesis via the canonical BKDH-dependent pathway. On the other hand, in certain environments, such as polymicrobial infections, organisms can compete for nutrients used in shared metabolic pathways, as is the case of amino acids. For instance, BCAAs can become scarce which can affect BCFAs synthesis via BKDH. Catabolism of BCAAs via the Buk-Ptb pathway in the acyl-CoA-forming direction would result in the release of volatile compounds into the polymicrobial environment (i.e. BCCAs, BC-aldehydes). These in turn would be available for BCFAs synthesis via MbcS. Our data suggest that MbcS is particularly important during polymicrobial infections where it helps *S. aureus* to establish a competitive relationship with other microbial species. Our laboratory is actively working to understand the role of MbcS in polymicrobial environments, including during co-infection.

**Figure 7.**
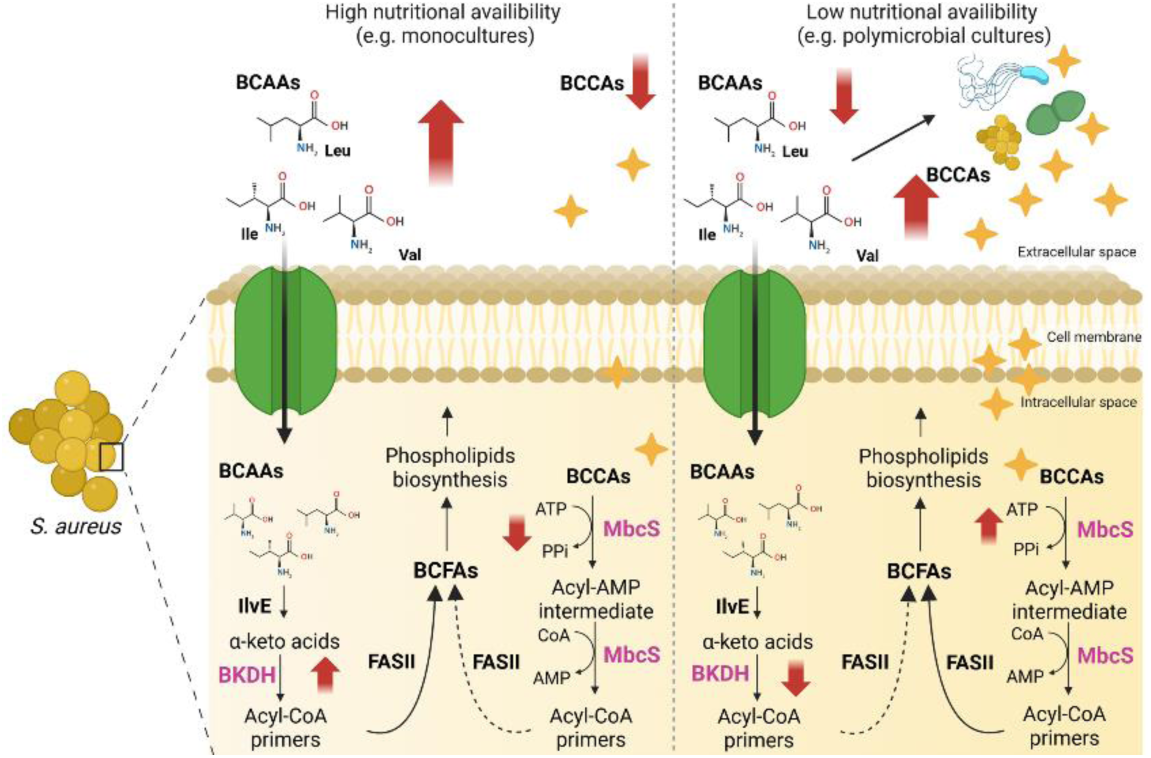
Proposed mechanism by which *S. aureus* uses the MbcS-dependent pathway for BCFAs synthesis during nutrient limitation. Two scenarios are depicted – a nutrient replete condition (left) and a nutrient-depleted condition (right). IlvE, BCAA aminotransferase; FASII, type II fatty acid synthase.

## Materials and Methods

### Bacterial strains, growth media, and culture conditions

Strains used in the present study are listed in **Table S1**. To routinely cultivate *S. aureus* strains, tryptic soy both (TSB) containing 0.25% (wt/vol) dextrose (BD Biosciences) or complete chemically defined medium (CDM, pH 6.5) (89) were used. Briefly, CDM medium was formulated with the L-form amino acids alanine (672 μM), arginine (287 μM), aspartic acid (684 μM), cysteine (166 μM), glutamic acid (680 μM), glycine (670 μM), histidine (129 μM), isoleucine (228 μM), leucine (684 μM), lysine (342 μM), methionine (20 μM), phenylalanine (240 μM), proline (690 μM), serine (285 μM), threonine (260 μM), tryptophan (50 μM), tyrosine (275 μM), valine (684 μM); the vitamins thiamine (56 μM), nicotinic acid (10 μM), biotin (0.04 μM), pantothenic acid (2.3 μM); and the small molecules MgCl_2_ (1000 μM), CaCl_2_ (100 μM), monopotassium phosphate (40,000 μM), dipotassium phosphate (14,700 μM), sodium citrate dehydrate (1400 μM), magnesium sulfate (400 μM), ammonium sulfate (7600 μM), and glucose (27,753 μM). For growth with high concentrations of branched-chain carboxylic acids (BCCAs), 2 g L^−1^ sodium bicarbonate was added to CDM to increase buffer capacity. Blood agar plates were used to propagate the *lpdA::kan^+^ mbcS::erm^+^* double mutant and its derivatives to mitigate the severe growth defect on TSB. *Escherichia coli* strains were grown in lysogeny broth (LB) medium without glucose (10 g L^−1^ tryptone, 5 g L^−1^ yeast extract, and 10 g L^−1^ sodium chloride) (90). When necessary, media were solidified with agar at 1.5% [w/v] and supplemented with antibiotics at the following concentrations to maintain selection: ampicillin (Ap) 100 μg mL^−1^, chloramphenicol (Cm) 10 μg mL^−1^, erythromycin (Erm) 10 μg mL^−1^, trimethoprim (Tm) 10 μg mL^−1^, or tetracycline (Tc) 1.5 μg mL^−1^. When required, media were supplemented with the short BCCAs *i*C_4_, *i*C_5_, and *a*C_5_ (Sigma-Aldrich) to the indicated final concentrations or with the BCFA *a*17:0 (Sigma-Aldrich) to a final concentration of 0.5 mM. Anhydrotetracycline (aTc) was used to induce the expression of genes under the control of the P*_tet_* promoter when indicated. Unless otherwise noted, all strains were grown at 37°C. A single dilution scheme was used to grow cells for genetic complementation and competition assays. Briefly, after streaking from frozen stocks, single colonies were used to initiate overnight cultures into 16×125 mm borosilicate glass tubes containing 2 ml of either plain TSB or TSB supplemented with *a*17:0, with rotation. Next day, overnight cultures were diluted 1:100 or 1:50 into 16×125 mm glass tubes with 4 ml fresh TSB and re-grown to exponential phase (up to 4 h for some strains). Cells were collected at an OD_600_ of 0.8 ± 0.2 and pellets were washed twice with plain CDM. Samples were inoculated into either plain CDM or CDM supplemented with the short BCCAs *i*C_4_, *i*C_5_, and *a*C_5_ in a 96-well plate to an OD_600_ of 0.05. Plates were incubated overnight in a microtiter plate shaker incubator (Stuart SI505) at 600 rpm and cell density measured in a computer-controlled Synergy H1 plate reader (BioTek/Agilent) running Gen5 software ver3.14.

### Genetic techniques

Oligonucleotides used in this study were synthesized by Integrated DNA Technologies (IDT; Coralville, IA) and are listed in **Table S2**. Plasmids used in this study are listed in **Table S3** and were constructed using Gibson assembly as described previously (91) or restriction enzyme-dependent methods. All molecular biology enzymes and a PCR product purification kit were purchased from New England Biolabs. A plasmid purification kit was purchased from Zymo Research. *E. coli* DH5α was used as host for plasmid constructions; plasmids were then transferred into *S. aureus* strain RN4220 by electroporation as previously described (92). Plasmids and marked mutations were moved between *S. aureus* strains via Φ85-mediated transduction (93). All plasmids were verified by whole plasmid sequencing (Plasmidsaurus), and all chromosomal alleles and integrated plasmids were verified by PCR and/or Sanger sequencing (Azenta).

### Phylogenetic analysis

All assembled Staphylococcaceae genomes with predicted proteins were downloaded from the NCBI database on 06/24/2025 (https://www.ncbi.nlm.nih.gov/datasets/genome/?taxon=90964). PhyloPhlAn v3.1.6B (94) was used to construct a phylogenetic tree of the downloaded genome assemblies. The “s *Staphylococcus_aureus*” marker gene set from PhyloPhlAn was used to build the phylogeny, with the program configured (using the ‘phylophlan_write_config_file’ command) with the following parameters: “-d a --force_nucleotides --db_aa diamond --map_aa diamond --map_dna diamond --msa mafft –trim trimal --tree1 fasttree --tree2 raxml”, and run (using the ‘phylophlan’ command) using the following parameters “-t a --diversity medium --fast - -force_nucleotides”. The functional annotations provided by NCBI for each protein were searched for five target genes of interest (*buk*, *ptb*, *betA*, *betB*, and *mbcS*). A case insensitive text search was conducted across all downloaded proteins, with Buk proteins identified as having a description of “butyrate kinase”, Ptb proteins as having a description of “butyryltransferase”, BetA proteins as having a gene symbol of “*betA*”, and BetB proteins as having a gene symbol of “*betB*”. MbcS proteins were identified by their description and proximity along their respective genomes to *betB*; A protein was considered a putative *mbcS* if it (1) had a functional description of “acyl-CoA synthetase” or “AMP-binding enzyme”, or a gene symbol of “*mbcS*”, and (2) it was ≤5 genes downstream of a *betB* gene. The presence or absence of the five target genes within each of the genomes was visualized using GraPhlAn v1.1.3 (https://github.com/biobakery/graphlan). A phylogenetic analysis of all *mbcS* and *mbcS*-like proteins was conducted to assess the evolutionary context of *mbcS* and to identify additional *mbcS* genes that were missed in our identification approach (i.e., were not correctly described functionally or were not in close genomic proximity to *betB*). All proteins with the functional description of “acyl-CoA synthetase” and “AMP-binding”, or the gene symbol of “*mbcS*” or “*acs*” were retrieved from the downloaded genomes. These proteins were aligned using mafft v7.453 (“--auto”) (94), with the resulting alignment used by iqtree v3.0.1 (“-m ‘LG+I+G4’; --fast”, best model selected using ModelFinder (95, 96) for phylogenetic inference. The resulting phylogeny was midpoint rooted and visualized using GraPhlAn. The known MbcS proteins (i.e., correct description and proximity to *betB*) were used to identify a putative MbcS protein clade, which contained many sequences unidentified by our first analysis.

### Plasmid construction

#### Construction of complementation plasmids

##### pCS02

The native promoter of the operon containing the *codY^+^* gene (358-bp sequence upstream of the *xerC* gene) and the 774-bp fragment containing the *S. aureus codY^+^*coding sequence were amplified from pKM25 with the forward primer containing a NotI cutting site and the reverse primer containing a BamHI cutting site. Both the vector backbone, pKK22, and the *codY^+^* insert were digested with BamHI and NotI and the resulting digested products were cleaned up using a PCR purification kit. The purified vector and insert were ligated using T4 DNA ligase and the resulting plasmid was then transformed into *E. coli* DH5a.

##### pMF020

An 894-bp fragment containing *ptb^+^* and a 1089-bp fragment containing *buk^+^* coding sequences of each gene from *S. pseudintermedius* were codon-optimized for *S. aureus*, synthesized, and cloned into pWY54 by GenScript. The resulting plasmid was then transformed into *E. coli* DH5α. This places the production of *Spptb^+^Spbuk^+^*under the control of the inducible P*_tet_* promoter.

##### pMF022

A gBlock was used to generate a 1584-bp fragment containing the MUA81_01165*^+^* coding sequence from *S. simulans* strain IVB6244 genome (GenBank accession number CP094697). Plasmid pWY54 was amplified with primer pair oMF233/oMF234. The DNA fragments were assembled by the Gibson method, and the DNA mixture was transformed into *E. coli* DH5α, resulting in pMF022. This places the production of MUA81_01165 under the control of the inducible P*_tet_* promoter and the *S. aureus mbcS* Shine Dalgarno sequence.

#### Construction of overproduction plasmids

##### pMF017 (for *SpPtb* production)

An 894-bp fragment containing the *ptb^+^* coding sequence was amplified from *S. pseudintermedius* using primer pair oMF208/oMF209. In parallel, pTEV5 (97) was amplified using primer pair oMF210/oMF211. The DNA fragments were assembled by the Gibson method, and the DNA mixture was transformed into *E. coli* DH5α, resulting in pMF017.

##### pMF018 (for *SpBuk* production)

A 1059-bp fragment containing the *buk^+^* coding sequence was amplified from *S. pseudintermedius* using primer pair oMF212/oMF213. In parallel, pTEV5 (97) was amplified using primer pair oMF214/oMF215. The DNA fragments were assembled by the Gibson method, and the DNA mixture was transformed into *E. coli* DH5α, resulting in pMF018.

### GFP reporter assay

Single colonies of cells carrying the indicated reporter fusion were inoculated into sterile borosilicate glass tubes (16 × 125 mm) containing 2 mL of TSB supplemented with Cm 10 μg mL^−1^ to maintain selection. Cells were grown to a stationary phase overnight. Cells were pelleted, washed twice with phosphate-buffered saline (PBS), and resuspended in PBS. 100 μL of the cell suspension was transferred into a flat bottom black 96-well plate (Corning). Fluorescence was measured using a Synergy H1 plate reader (BioTek/Agilent) by tuning the monochromator to 485 nm and 535 nm (excitation/emission, respectively). Relative fluorescence units (RFUs) were calculated by subtracting the fluorescence of plain PBS and dividing by OD_600_ to correct for cell density.

### Competition experiments

From the single dilution scheme described above, exponential-phase cells were diluted in TSB to an OD_600_ of 0.05 (∼10^6^ colony forming units (CFU) ml^-1^). From this initial culture, cells were inoculated to ∼10^5^ CFU in 1 ml of fresh TSB in 5 ml conical tubes for the respective monocultures and co-cultures. 5 ng mL^−1^ aTc was added to the cultures to induce expression of p*SambcS*^+^. For co-cultures, differentially marked strains were mixed at 1:1 ratio (i.e. ∼10^5^ CFU of each strain in 1 ml). Initial cultures were spotted on tryptic soy agar (TSA) containing antibiotics to selectively count CFUs. For competitions, cells were grown to stationary phase overnight with shaking, followed by spotting TSA plates containing appropriate antibiotics for selective CFU counting. Competition indices were calculated as previously described (61, 62). Calculated indices, which evaluate mutant over WT ratios, were compared to 1, at which index there was no competition.

### Protein production

#### *Sp*Ptb

Plasmid pMF017 was transformed into *E. coli* BL21 (DE3). The resulting strain was grown in 10 ml LB supplemented with 100 μg ml^−1^ Ap at 37°C in 50 ml conical tubes. Overnight cultures were subcultured at a 1:100 dilution into 500 mL of LB supplemented with 100 μg ml^−1^ Ap in a 2 L flask. Cells were grown at 37°C with shaking at 200 RPM in a Brunswick Innova 42R Stackable Incubator Shaker (Eppendorf) until an OD_600_ of ∼0.4. Expression was induced using 0.5 mM isopropyl β-D-1-thiogalactopyranoside (IPTG), followed by a 4 h incubation at 25°C with shaking at 200 RPM. Cells were harvested by centrifugation at 7,000×g for 10 min, and the resulting pellets were stored at −80°C until purification.

#### *Sp*Buk

Plasmid pMF018 was transformed into *E. coli* BL21 (DE3) cells. The resulting strain was grown in 10 ml LB supplemented with 100 μg ml^−1^ Ap at 37°C in 50 ml conical tubes. Overnight cultures were subcultured at a 1:100 dilution into 500 mL of LB medium supplemented with 100 μg ml^−1^ Ap in a 2 L flask. Cells grew at 37°C with shaking at 200 RPM in a Brunswick Innova 42R Stackable Incubator Shaker (Eppendorf) reaching an OD_600_ of ∼0.4. Expression was induced using 0.5 mM IPTG, followed by a 6 h incubation at 18°C with shaking. Cells were harvested by centrifugation at 7,000×g for 10 min, and the resulting pellets were stored at −80°C until purification.

### Protein purification

Pellets were resuspended in 20 mL of buffer 1 (50 mM Tris–HCl pH 8.0, 300 mM NaCl, 10 mM imidazole) containing 1 mg mL^−1^ lysozyme (Thermo Scientific). Cells were lysed using a Digital Sonifier (Branson) with 2-sec pulses (25% amplitude) separated by 5-sec pauses on wet ice for 2- to 3 minutes. Cellular debris were removed by centrifugation at 12,000×g for 15 min at 4°C. The proteins, fused to a TEV-cleavable *N*-terminal hexahistidine (His_6_) tag, were purified by Ni^2+^ affinity chromatography. Briefly, the lysate was loaded on a 5 mL bed of nickel-nitrilotriacetic acid resin (Thermo Scientific) pre-equilibrated with buffer 1. The column was then washed with six column volumes of buffer 2 (50 mM Tris–HCl pH 8.0, 300 mM NaCl, 20 mM imidazole). The proteins were eluted with six column volumes of buffer 3 (50 mM Tris–HCl pH 8.0, 300 mM NaCl, 250 mM imidazole). Eluted protein fractions were analyzed using SDS-PAGE, and those fractions containing the target protein were pooled. His-tagged proteins were concentrated using 15 ml Amicon Ultra centrifugal tubes with a 10 kDa molecular weight cut-off (Sigma), and buffer exchanged sequentially to buffer 4 (50 mM Tris–HCl pH 7.5, 250 mM NaCl, 0.5 mM EDTA), storage buffer 5 for *Sp*Ptb (50 mM Tris–HCl pH 7.5, 250 mM NaCl, 10% glycerol) and storage buffer 6 for *Sp*Buk [50 mM Tris–HCl pH 7.5, 200 mM NaCl, 10% glycerol, 5 mM dithiothreitol (DTT)]. Protein concentrations were determined by measuring absorbance at 280 nm (A280) using a NanoDrop 2000c spectrophotometer (Thermo Scientific) and purity was evaluated using SDS-PAGE. Due to difficulties encountered during His_6_ tag cleavage with TEV protease, tagged proteins were used for in vitro studies.

### Hydroxamate enzyme assay

*Sp*Buk activity was assessed through the colorimetric hydroxamate assay with modifications (37, 56, 57). Briefly, purified His_6_-*Sp*Buk was mixed with 2 µmol ATP, and 80 µmol neutralized hydroxylamine hydrochloride in buffer (50 mM HEPES, pH 7.5; 5 mM MgCl_2_; 5 mM tris (2-carboxyethyl) phosphine (TCEP)). The enzymatic reaction was performed in a 96-well plate at 37°C and initiated by the addition of various amounts of short-chain carboxylic acids in a final volume of 200 µl. Carboxylic acids were purchased from Sigma and were of the highest quality possible (99% purity).The reaction was stopped after 1 h by the addition of 40 µl of 50% trichloroacetic acid (TCA). Precipitated protein was removed by centrifugation at 2000 x *g* for 5 min. The supernatant was transferred to a 1.5 ml centrifuge tube and color was developed by the addition of 400 µl of 1.25% ferric chloride solution in 1 M hydrochloric acid (HCl). Formation of acyl-hydroxamate was measured using absorbance at 540 nm (A_540_) in a Synergy H1 plate reader (BioTek/Agilent)

### Liquid chromatography-mass spectrometry (LC–MS)

#### (i) Reagents and chemicals

All reagents and chemicals used were LC-MS (Optima) grade. Water, acetonitrile, isopropanol, and methanol were purchased from Honeywell, and formic acid was purchased from Thermo-Scientific. 3-Hydroxypropyl-mercapturic acid-d3 (3-HPMA-d3) and Isobutyryl-Coenzyme A were purchased from Cayman chemicals.

#### (ii) Preparation of standards and internal standard stock solutions

Coenzyme A was resuspended in 660 µl of HEPES buffer pH 7.5 to generate a 3.83 mg ml^-1^ stock solution (stock A). A 1 mg mL^-1^ solution of Isobutyryl-Coenzyme A (IB-CoA) was prepared by dissolving IB-CoA in HEPES buffer pH 7.5 (stock B). Next, stock A and stock B were diluted in extraction buffer (1 mg mL^-1^ solution of the 3-HPMA-d3 was prepared in methanol and diluted to 1 µg mL^-1^ to make extraction buffer) to generate stock C (20 µg ml^-1^ of CoA and 10 µg ml^-1^ IB-CoA). Stock C was then serially diluted to generate calibration curve standards ranging from 20 ng ml^-1^ to 10 µg ml^-1^ (IB-CoA) and 40 ng ml^-1^ to 20 µg ml^-1^ (CoA).

#### (iii) Sample preparation

Buk and Ptb activities were assessed via LC-MS. Reactions were done in triplicate and the reaction mixture consisted of 200 mM HEPES pH 7.5, 2 µmol MgCl_2_, 2 µmol ATP, 1 µmol TCEP, 98.6 nmol CoA, 125 nM Buk, 1.25 µM Ptb, and 10 mM 2 µmol isobutyric acid. Both Buk and Ptb were denatured by incubating enzyme at 95°C for 10 minutes and used in reactions as negative controls to show the reaction was enzyme-dependent. The reactions were initiated by adding isobutyric acid and incubated at 37°C for 2 hours. Tubes containing the reaction mixture were then placed on ice. For IB-CoA measurements, to the 40 µl of samples was added 360 µl of extraction buffer and vortexed. For CoA measurements, the samples prepared for IB-CoA measurements were diluted 2.5-fold using extraction buffer. The samples were transferred to the MS vials for data acquisition. Blanks (solvent alone) were injected on either side of test samples to assess sample carryover. Data were processed using Target Lynx 4.1. The relative quantification values of analytes were determined by calculating the ratio of peak areas of transitions of samples normalized to the peak area of the internal standard.

#### (iv) LC-MS methods

This method is designed to measure CoA and Isobutyryl-CoA by UPLC-MS system. The samples were resolved on a Acquity UPLC HSS T3, 1.8 µm, 2.1 x 100 mm column online with a triple quadrupole mass spectrometer (Xevo-TQ-S, Waters Corporation, USA) operating in the multiple reaction monitoring (MRM) mode.

The LC gradient method started with 98% of mobile phase A (10 mM ammonium formate in water) that involved a gradient change from 2% B (ACN:water [95:5] with 10 mM ammonium formate) to 100% phase B in 2.0 minutes after an initial lag phase of 0.5 minutes with flow rate 0.6 ml/min. The column was maintained at 40 °C and injection volume was kept at 5 µL. The autosampler was maintained at 15 °C. The column eluent was introduced directly into the TQS mass spectrometer by electrospray operating in positive mode at a capillary voltage of 3.0 kV and a sampling cone voltage of 2 V. The desolvation gas flow was set to 1000 l/h and the desolvation temperature was set to 500 °C. The cone gas flow was 150 l/h and the source temperature was set to 150 °C. The sample cone voltage and collision energies were optimized for the analyte to obtain maximum ion intensity for parent and daughter ions using “IntelliStart” feature of MassLynx software (Waters Corporation, USA). The instrument parameters were optimized to gain maximum specificity and sensitivity of ionization for the parent and daughter ions. Signal intensities from all MRM Q1/Q3 ion pairs for the analyte were ranked to ensure selection of the most intense precursor and fragment ion pair for MRM-based quantitation. This approach resulted in selection of cone voltages and collision energies that maximized the generation of each fragment ion species. The details for the calibration curves for standards are included in the *Supplementary Materials* section (**Table S4**). CoA was detectable as low as 20 ng ml^-1^ (limit of detection [LOD]) and showed 40 ng ml^-1^ as the lowest limit of quantification (LLOQ). While IB-CoA showed the LOD and LLOQ as 10 ng ml^-1^ and 20 ng ml^-1^, respectively. The quantification range (QR) for CoA and IB-CoA were 40 ng ml^-1^ to 20 µg ml^-1^ and 20 ng ml^-1^ to 10 µg ml^-1^, respectively. The relative quantification values of analytes were determined by calculating the ratio of peak areas of transitions of samples normalized to the peak area of the internal standard.

### Statistical analysis

Prism version 9.0 (GraphPad Software) was used to perform all statistical analyses. Statistical significance was calculated using measurements from at least three biological replicates and one-way analysis of variance (ANOVA). If significant, analyses were followed by Tukey’s honestly significant difference (Tukey’s HSD) tests for pairwise comparisons. For those data that did not follow a normal distribution pattern, significance was calculated using Kruskal-Wallis test followed by Dunn’s multiple comparison test. The statistical significance of competitive indexes was calculated using the one sample Wilcoxon test, where the hypothetical value was 1 (i.e., no competition). Statistical significance was assumed at a *p* value of <0.05.

## Author Contributions

SRB conceived the study; MCDSF and SRB conceptualized the research goals and aims; MCDSF and TGS performed the investigations and analyzed the data, SRB, MCDSF, and TGS prepared the original manuscript draft; SRB and MCDSF prepared the final manuscript.

## Acknowledgements

We thank Dr. Lindsey Shaw and Mary-Elizabeth Jobson at University of South Florida for the generous gift of the *S. aureus* transcription factor mutant library. We thank Dr. Jessica Gilbertie and Douglas Pluta at Virginia Tech for the gift of the *Staphylococcus pseudintermedius* clinical isolate. We thank Dr. Amrita Cheema, Dr. Shivani Bansal, and Satinder Kaur in the Georgetown University Metabolomics and Proteomics Shared Resource for help with LC-MS analysis, which is partially supported by NIH/NCI/CCSG grant P30 CA051008-29S2. We thank Dr. Carl Stone and Dr. Dennis DiMaggio for their technical expertise in pCS02 construction. We also thank members of the Georgetown University Microbial Interest Group, Brinsmade Lab members, and Dr. Jeffrey Boyd for their helpful comments and discussions.

